# Patrolling human SLE haematopoietic progenitors demonstrate enhanced extramedullary colonisation; implications for peripheral tissue injury

**DOI:** 10.1101/2021.01.15.426761

**Authors:** Ioannis Kokkinopoulos, Aggelos Banos, Maria Grigoriou, Anastasia Filia, Theodora Manolakou, Themis Alissafi, Nikolaos Malissovas, Ioannis Mitroulis, Panayotis Verginis, Dimitrios T. Boumpas

## Abstract

Systemic Lupus erythematosus (SLE) is an autoimmune disease where bone-marrow-derived haematopoietic cells have a key role in its pathogenesis with accumulating evidence suggesting an aberrant function of haematopoietic stem/progenitor cells (HSPCs). By employing next-generation sequencing, we compared the gene transcription signatures of CD34^+^ HSPCs deriving from either the bone marrow or HSPCs patrolling the bloodstream of healthy and individuals with SLE, seeking common transcriptional pathways that may have been modified between steady and disease states. Our findings indicate that circulating and bone marrow-derived HSPCs are distinct in steady and diseased states. Non-mobilised, SLE-derived circulating HSPCs demonstrated enhanced engrafting and altered differentiation capacities. Importantly, xenotransplantation of circulating HSPCs in humanised mice showed that human peripheral blood HSPCs possess the ability for extramedullary organ colonisation to the kidneys. SLE CD34^+^ HSPCs homing and engraftment at extramedullary sites such as the spleen and kidneys may participate in peripheral tissue injury.

## INTRODUCTION

Long-term hematopoietic stem cells (LT-HSC) differentiate through short-term HSC (ST-HSC) and then multipotent haematopoietic progenitor (MPP) stages into lineage-restricted progenitors such as lymphoid, myeloid, or megakaryocyte/erythroid progenitors. HSCs are an integral part of the immune response with the ability to sense inflammatory stimuli in infectious and chronic inflammatory diseases. Importantly, prolonged exposure to inflammatory stimuli during chronic inflammatory diseases has long-lasting effects on the Bone Marrow (BM) cell output’s nature through epigenetic modifications in hematopoietic stem and progenitor cells (HSPCs) (1–5). Dysregulation of HSPC activity in the BM has been reported in several chronic inflammatory diseases, including inflammatory bowel disease, experimental spondyloarthritis, atherosclerosis and systemic lupus erythematosus (SLE) (6–9).

SLE is the prototypic systemic autoimmune disease characterized by inflammation and damage in several organs and a disease course, where periods of treatment-induced remissions alternate with flares. In SLE, most cells participating in the pathogenesis of SLE originate from BM HSPCs. Murine and human SLE HSPC’s gene expression program is biased towards myelopoiesis and GMP progenitors (10, 11). Based on these findings we reasoned that in lupus, systemic inflammation may enhance both medullary and extramedullary myelopoiesis to meet the increased demand for effector cells in the periphery. In this setting, HSCPs may emigrate from the bone marrow to peripheral tissues and seed the red pulp of the spleen or other inflamed tissues to produce myeloid cells (neutrophils and monocytes) and contribute to peripheral pathology (11).

Herein, we investigate how HSC in the BM and patrolling HSPCs in the circulation respond to lupus systemic inflammatory environment (12). To this end, we collected BM and PB samples from healthy subjects and SLE patients and performed mRNA-seq in isolated human CD34^+^ HSPCs. Using HSC human surface markers, we confirm the presence of HSC in the PB, with the fast-cycling MPPs being increased in the PB of SLE individuals. Our analysis also indicates that PB- and BM-derived CD34^+^ progenitors have distinct gene expression signatures in transcriptional networks and cellular functions both in healthy and SLE patients, as well as different migratory and metabolic attitudes, between them. We also identify that the histone deacetylase *SIRT7* that has recently been shown to be relevant to HSC rejuvenation and reconstitution (13–15), is highly expressed in PB-HSCs in comparison to their BM-counterparts. Using adult humanised mice as hosts, we report that human PB SLE CD34^+^ and MPP cells show enhanced homing and extramedullary colonisation in humanised mice compared to Healthy PB counterparts. SLE CD34^+^ HSPCs homing and engraftment at extramedullary sites such as the spleen and kidneys may participate in local pathology.

## RESULTS

### Human SLE PB HSPCs show an enhanced extramedullary gene expression profile

SLE involves the dysregulation of the HSCs, resulting in a multi-organ disease phenotype (10) that could potentially involve circulating patrolling progenitors. To this end, we isolated total RNA from magnetically-isolated CD34^+^ cells extracted from the periphery (PBMCs) and the bone marrow aspirates [bone marrow mononuclear cells (BMMCs)] of age-matched healthy and SLE individuals and subjected them to NGS mRNA-seq.

CLINVAR database for human disease variants (16, 17) analysis on differential gene expression (DGE) indicated that SLE haematopoietic circulating progenitors differentially expressed key genes involved in migration and extramedullary colonisation (*IFN-γ, IL-8R, CCL3L1-CCL3*) and SLE-specific immune regulation (*CTLA4, STAT4, PCD1*), when compared to Healthy PB circulating progenitors (Figure **1A**). REACTOME Reactions and Pathways analyses (18) revealed significant changes in gene expression involved in extracellular matrix (ECM) organisation, immune system activation through CD3, and cytokine/chemokine regulation that involved IL-10 signalling and trafficking in inflammation sites (12, 19), with the majority of genes in the SLE-derived PB progenitors, being upregulated in comparison to Healthy-derived PB (Figure **1B**). Finally, KEGG pathways analysis (20) was indicative of DGE that involved the migratory and extramedullary increased potential (cytokine and cytokine-receptor interactions) as well as altered immune activation of circulating SLE progenitors when compared to their circulating Healthy progenitor counterparts (Figure **1C**).

**Figure 1.**
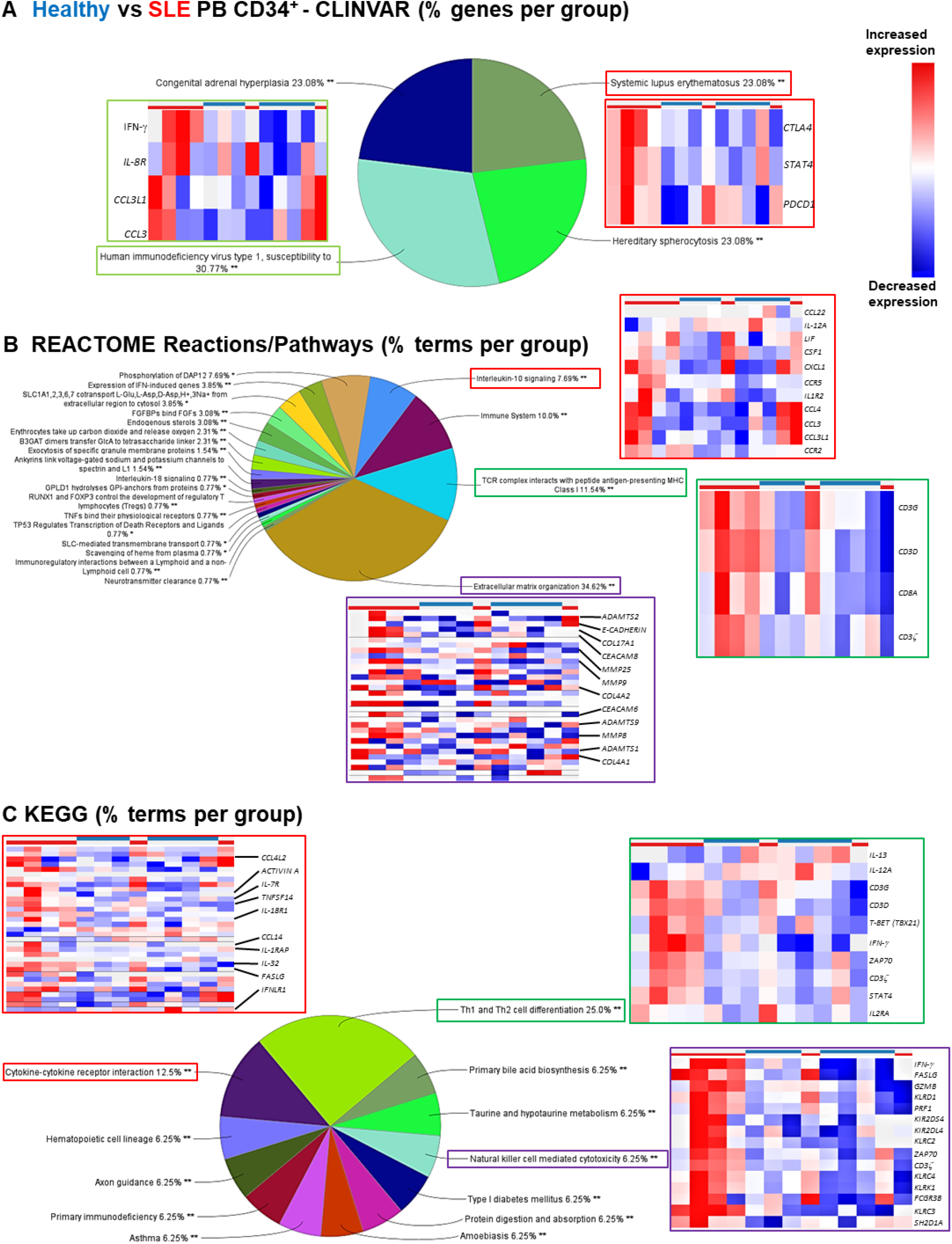
SLE PB CD34^+^ progenitors show upregulated expression of genes involved in extramedullary colonisation. (A) CLINVAR analysis of DE genes between Healthy PB (blue) vs SLE PB (red) CD34^+^ progenitors with relevant heatmaps of the bracketed GO terms. (B) REACTOME Reactions/Pathways analysis of DE genes between Healthy PB (blue) vs SLE PB (red) CD34^+^ progenitors with relevant heatmaps of the bracketed GO terms. (C) KEGG analysis of DE genes between Healthy PB (blue) vs SLE PB (red) CD34^+^ progenitors with relevant heatmaps of the bracketed GO terms. All analyses were performed with a p<0.05 with an FDR correction. *p<0.05, **p<0.01.

By employing the same analytical principle (CLINVAR, REACTOME Reactions/Pathways and KEGG) on DGEs between Healthy vs SLE BM CD34^+^ progenitors, although a similar immune reaction profile was evident, an extramedullary/migratory signature was not readily apparent in the SLE BM CD34^+^ HSPCs, as judged by cytokine and cytokine-receptor interaction gene expression profile (Supplementary Figure **1**). CLIVAR and KEGG analyses on DGE between Healthy PB vs BM and SLE PB vs BM CD34^+^ progenitors indicated that circulating progenitors (both Healthy and SLE) showed a decrease in oxidative phosphorylation gene expression. In contrast, Healthy BM progenitors had a non-migratory profile when compared to Healthy PB progenitors (Supplementary Figures **2A, B**).

Together, these results indicate that human lupus circulating HSPCs possess a transcriptomic signature that may promote an enhanced engraftment potential in relation to BM HSPCs.

### Human PB vs BM CD34^+^ HSPCs

In light of these differences, we next assessed whether the transcriptome is distinct among the four haematopoietic progenitor populations (PB SLE, PB Healthy, BM SLE and BM Healthy). Principal component analysis (PCA) depicting all the differentially expressed (DE) genes between those four groups, indicated distinct patterns in the transcriptome between BM- and PB-derived CD34^+^ progenitors, irrespective of disease status (four groups, Figures **2A, B**). This finding was also recapitulated when PB- and BM-derived progenitors were compared within the same disease setting (healthy or SLE, Figure **2C**).

**Figure 2.**
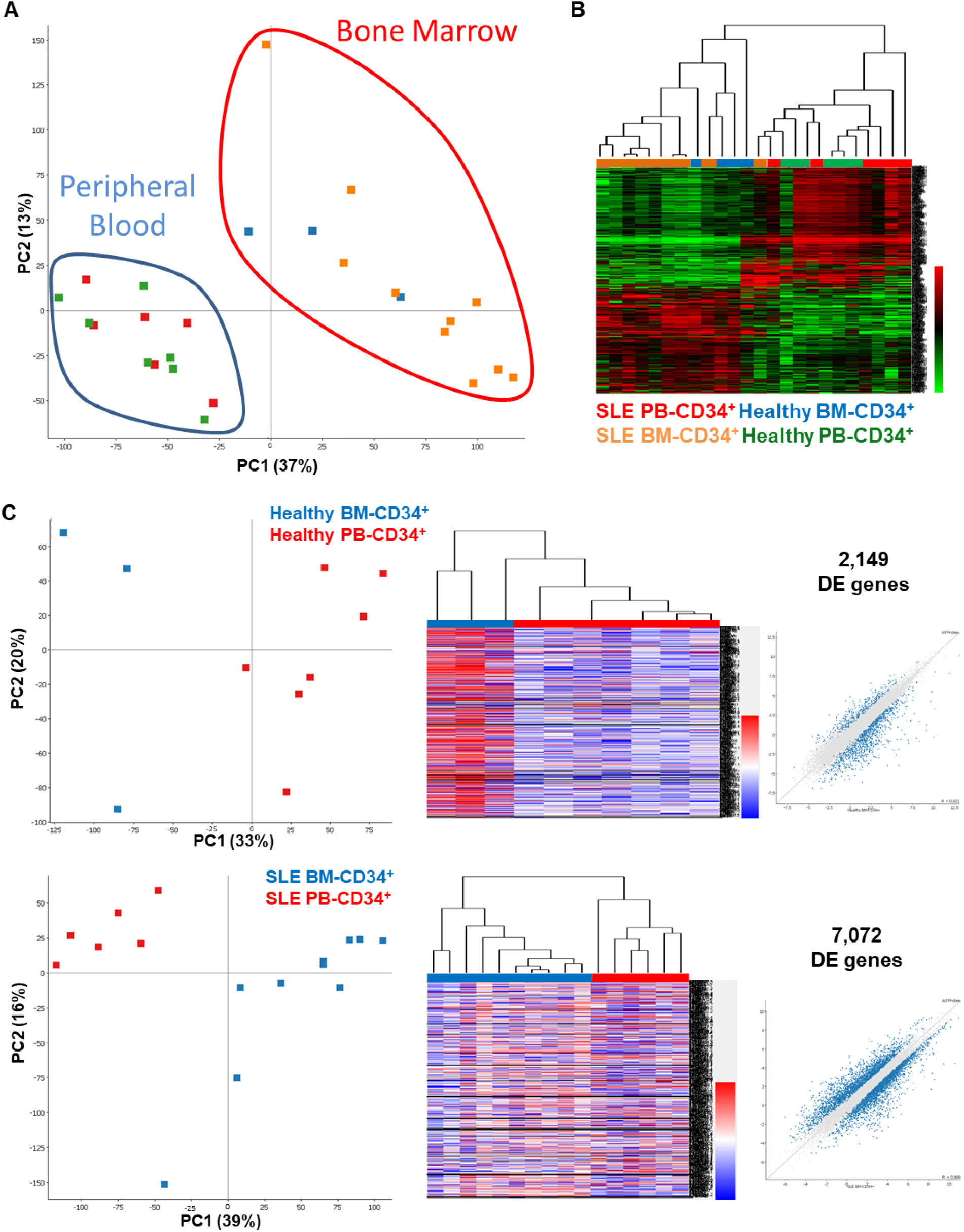
Circulating and niche haematopoietic progenitors have different transcriptomic signatures, irrespective of diseases status. (A) PCA analysis indicating the transcriptional clustering of human CD34^+^ progenitors from SLE PB (red squares, n=6), SLE BM (orange squares, n=10), Healthy BM (blue squares, n=3) and Healthy PB (green squares, n=7). (B) Heatmap representation of 12,659 DE genes depicting clustering of BM- and PB-derived haematopoietic progenitors in Healthy and SLE individuals. (C) PCA analysis, heatmap and scatter plots indicating the transcriptional clustering of human CD34^+^ progenitors from Healthy PB vs BM and SLE PB vs BM. EdgeR analysis indicated 2,149 DE genes between BM and PB CD34^+^ progenitors, while in SLE samples, 7,072 DE genes were detected between PB and BM.

Next, we set to explore if transcriptome analysis could reveal significant gene expression differences in CD34^+^ HSPCs, due to disease status (i.e. Healthy vs SLE). PCA analysis showed that PB (in contrast to BM) did not show strong segregation, bearing a smaller number of DE genes (Supplementary Figures **2C, D**).

We looked further into this comparison by comparing the DGE analysis of SLE DGE between PB and BM, by using the Healthy DGE PB vs BM output as a reference point. CLINVAR analysis confirmed an SLE-specific signature, providing a list of genes altered in PB SLE CD34^+^ HSPCs (Figure **3A**). Hierarchical clustering revealed that these genes were indeed differentially expressed in PB SLE versus PB Healthy progenitors. KEGG pathways analysis revealed that antigen presentation and processing that included Graft-versus-Host-Disease (GvHD), allograft rejection and oxidative phosphorylation, were overrepresented in SLE compared to healthy counterparts (Figure **3B**). Haematopoietic cell lineage, Th17 cell differentiation, antigen processing and presentation as well as cell cycle, were also enriched in the PB setting, with the majority of genes being elevated (except cell cycle GO) in the SLE PB-derived cell populations, in comparison to healthy PB-derived progenitors. These data pinpoint a potential discrepancy between SLE PB- and BM-derived CD34^+^ cells that may cause significant differences in progenitor engraftment and immune activation (21, 22). KEGG analysis of DE genes in SLE vs healthy CD34^+^ cells showed that genes involved in allograft rejection pathways were enriched in the SLE setting when compared to healthy CD34^+^ HSPCs (Figure **3B**).

**Figure 3.**
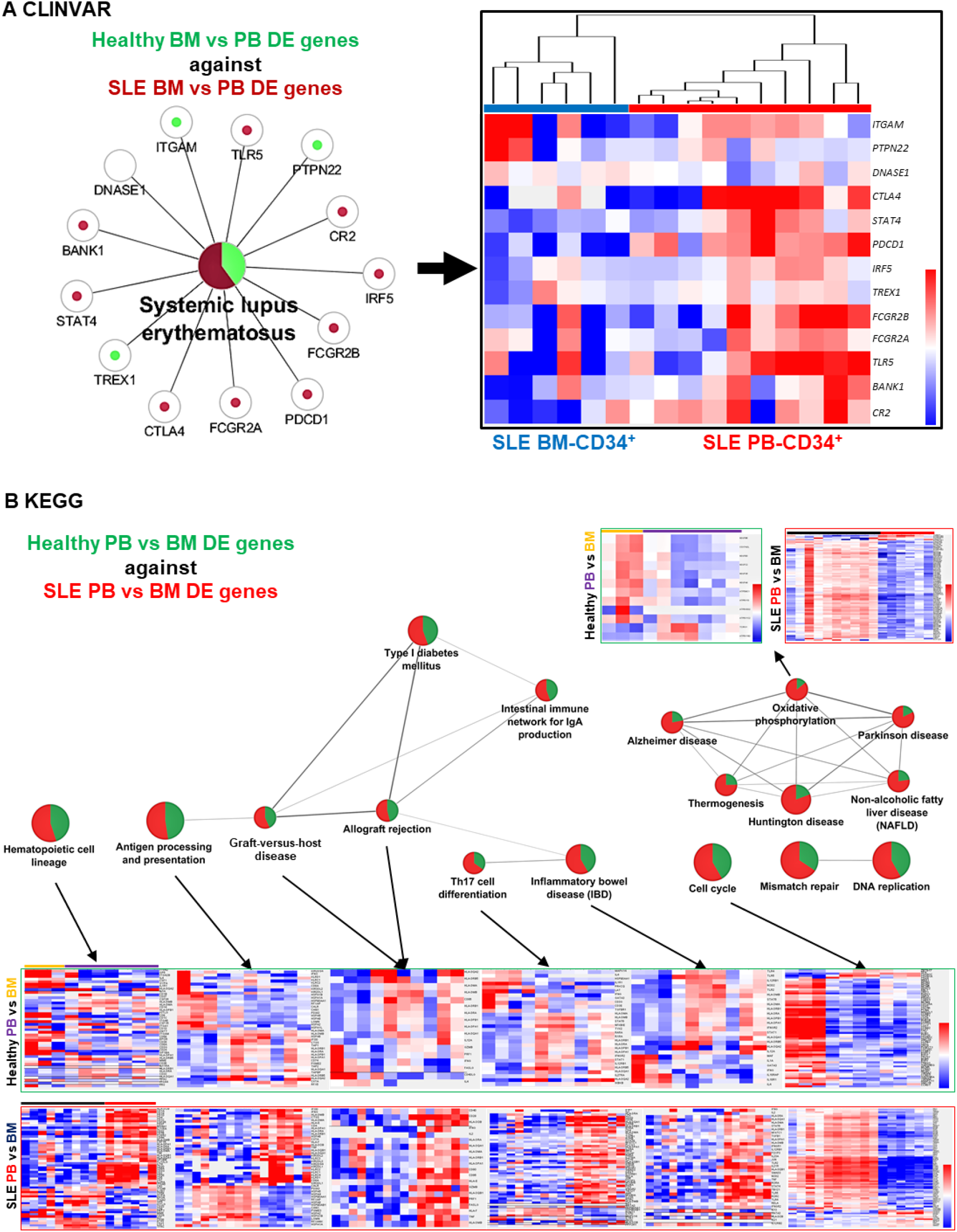
Different transcriptomic signatures in SLE PB and BM progenitors. (A) DE genes between SLE BM-derived and PB-derived progenitors were used as an input on CLINVAR database, using as a reference cut-off the DE genes found between healthy BM-derived and PB-derived progenitors. Graphical representation showing genes found to be either enriched in SLE (dark red) or Healthy (light green) progenitors, from genes reported in the CLINVAR database. A heatmap of these DE genes indeed indicated higher expression levels in SLE PB-derived CD34^+^, in comparison to SLE BM-derived CD34^+^. (B) KEGG pie chart analysis of DE genes between SLE-derived BM and PB CD34^+^progenitors (red), and Healthy BM vs PB-derived (green). Pie chart connecting lines indicate kappa-score relations. Heatmaps indicating DE genes that shown to be enriched in Healthy-derived PB (purple) when compared to Healthy-derived BM (light orange) progenitors, and SLE-derived PB (red) when compared to SLE-derived BM (black) progenitors. All analyses were performed with a p<0.05 with an FDR (Bonferroni and Heidelberg) correction.

Together, these analyses underscore the transcriptome differences in the circulating HSPCs in relation to BM CD34^+^ progenitors while underpinning their potential implication in SLE disease pathogenesis.

### Specific transcription factor binding sites predicted in PB- and BM-derived CD34^+^ progenitors

Next, we sought to identify potential Transcription Factor (TF) binding sites based on the DE genes identified between PB and BM, in both healthy and SLE patient samples. The analysis revealed a non-exhaustive list of potential TF sites which pinpointed unique TFs that may regulate a cohort of gene regulation (Figure **4**). SLE PB-derived progenitors showed a marked downregulation in *NRF1-dependent* DE genes (131 genes), while *LXR* (*NR1H3*)-dependent gene expression showed mixed DGE patterns (36 genes) (Figure **4A, B** and Supplementary Figure **3A**).

**Figure 4.**
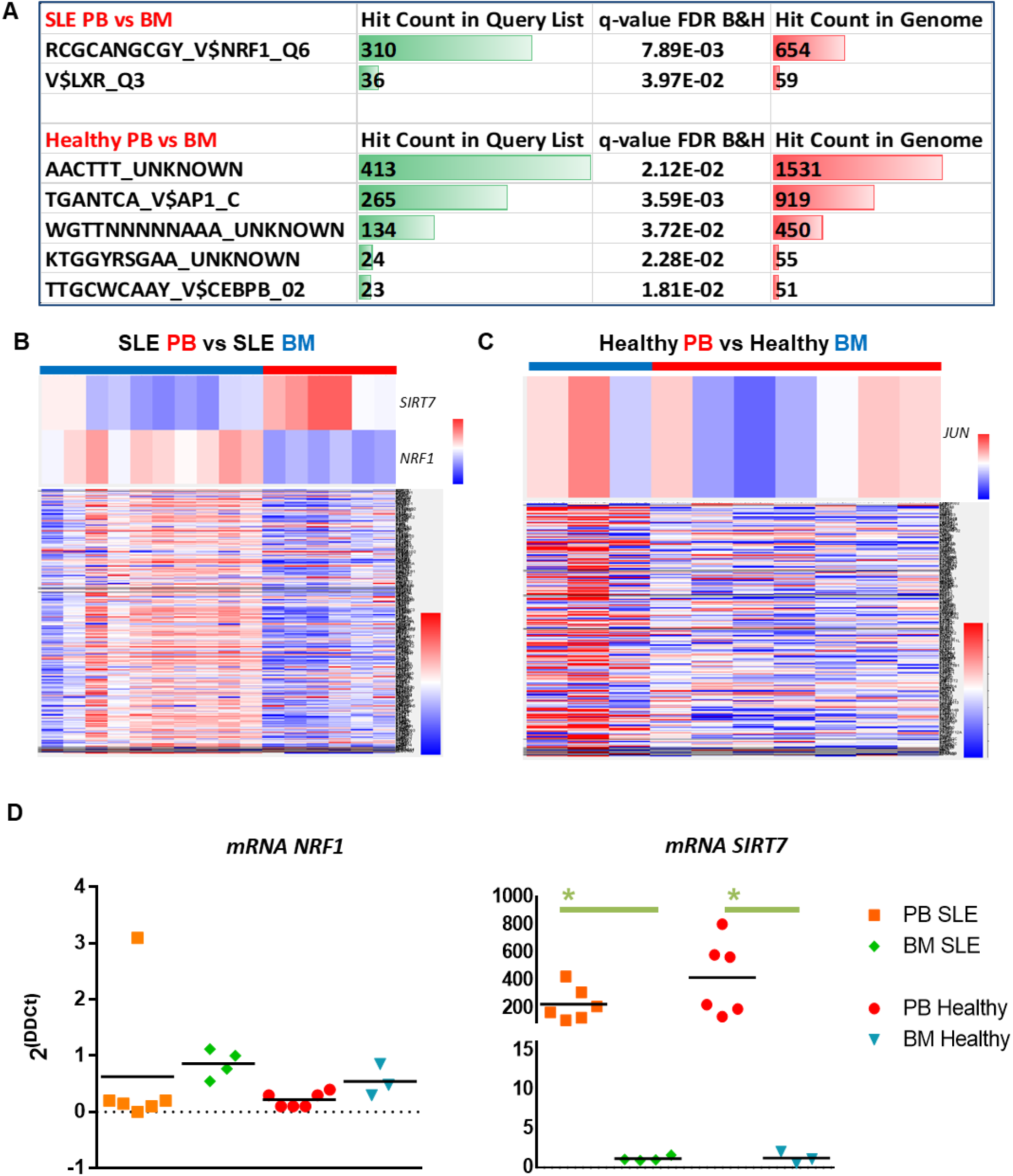
The NRF1-SIRT7 transcriptional signature is altered in PB CD34^+^ progenitors. (A) A table indicating the predicted TF binding sites (ToppgeneTM software) based on the DE gene input list. In the SLE PB vs BM comparison the NRF1 and LXR TFs were identified, while in the Healthy PB vs BM comparison, AP1 (JUN) and CEBPB were identified along those with unknown TF bindings site motifs. (B) Heatmaps of SIRT7 and NRF1 expression in SLE PB and SLE BM, along with 310 DE genes that are affected by NRF1 expression. (C) Heatmaps of JUN expression in Healthy PB and Healthy BM, along with 265 DE genes that are affected by JUN expression. (D) Real-time PCR graphs for NRF1 (ENST00000353868.5, ENST00000393230.6 variants) and SIRT7 (ENST00000575360.5, SIRT7-209, non-sense mediated decay) in all four groups. The expression of NRF1 was not statistically different in any of the four groups, albeit there was a trend of lower expression in the SLE PB. In contrast, SIRT7 expression was vastly and significantly elevated in PB CD34+ progenitors, in comparison to BM CD34^+^ progenitors. One-way ANOVA with Bonferroni’s post-hoc test, *p<0.05, n=3-6.

Due to the dynamic synergistic effect that *NRF1* induces on SIRT7 in relation to HSPCs (23, 24), several variants of both *NRF1* and *SIRT7* were explored (Figure **4D** and Supplementary Figure **3E**). *SIRT7* expression of its nonsense-mediated decay variant showed higher expression in PB, in relation to BM (both SLE and Healthy), yet with a two-fold decrease (non-statistically significant, but biologically important) in SLE PB, in comparison to Healthy PB. In healthy PB-derived progenitors, predicted expression of altered TFs *AP1* (*JUN*) - and *CEBPB*-dependent DE genes produced mixed differential gene expression patterns, (265 and 23 genes, respectively, Figure **4C** and Supplementary Figure **3B**).

We then investigated the prospect of TFs that may be related to the DE genes on PB- and BM-derived HSPCs between SLE and healthy patients. Prediction analysis showed a cascade of TF-dependent DE genes in the BM CD34^+^ progenitors, but none in the PB CD34^+^ progenitors (Supplementary Figure **3C, D**). In most of the predicted TFs, DE genes were downregulated in the SLE BM-derived progenitors compared to controls (data not shown). Yet, the TF family *E2F* and specifically *E2F1* (39 genes) revealed a cluster of DE genes that were upregulated in the SLE in comparison to healthy controls.

### SLE PB CD34^+^ progenitors showed an increased extramedullary differentiation in humanised mice

Based on the potentially altered migratory transcriptional profile of circulating HSPCs, we interrogated the behaviour of human PB CD34^+^ progenitors as xenotransplants. Human SLE or Healthy 2.5×10^5^ CD34^+^progenitors were injected into each of fourteen 2-3 month-old NBSGW mice (one human sample per mouse, Figures **5** and **6A**). Mice were sacrificed at 9, 11, 13, and 20-weeks post-injection to assess human cell colonisation potential. Mice did not develop alopecia, a characteristic of GvHD, up to 20-week post-injection (25). We observed enhanced engraftment kinetics and lymphocyte/myeloid differentiation potential from SLE PB CD34^+^ human cells when compared to healthy PB CD34^+^ cells, in the murine BM (Figure **5B**).

**Figure 5.**
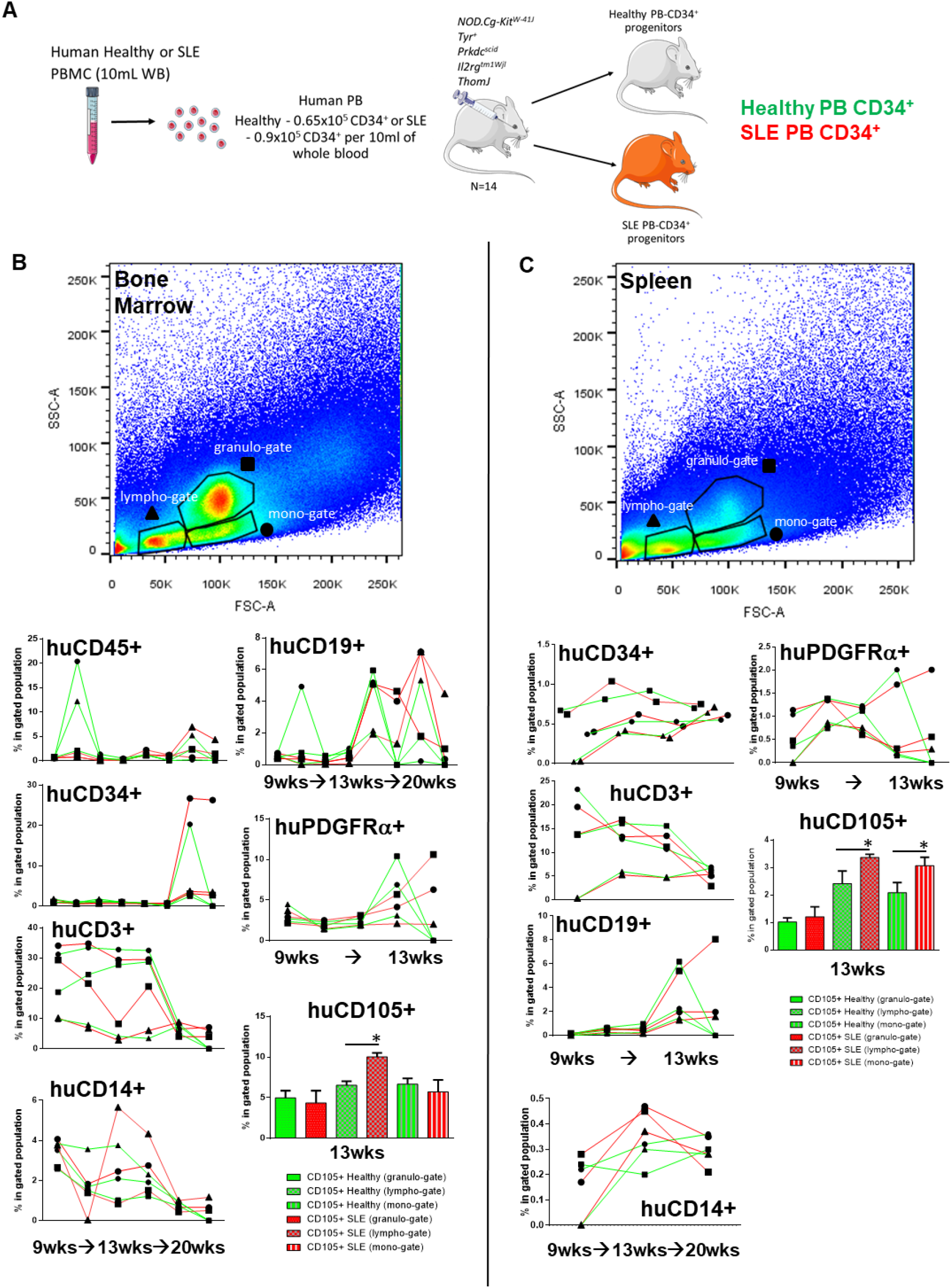
Altered kinetics of SLE CD34^+^ progenitors in BM and spleen. (A) Schematic representation of the xenotransplantation procedure of human PB CD34^+^ in 2-3-month-old mice, N=14. (B) Mice were sacrificed at designated time intervals and the BM-derived cells were subjected into immunostaining with antibodies against human-only surface markers, in order to assess human cell presence. (C) Mice were sacrificed at designated time intervals and the spleen-derived cells were subjected into immunostaining with antibodies against human-only surface markers, in order to assess human cell presence. Lympho-gates (triangle), granulo-gates (square) and mono-gates (circle) represent different immune cell subpopulations and therefore examined separately. Each triangle/dot/circle represents the average value for each time-point, and assigned gate.

In the spleen and kidneys differences in human CD19^+^, CD14^+^ and CD3^+^ cell percentages, between SLE- and Healthy-derived progenitors were evident (Figures **5C** and **6A**). Of interest, a substantial number of human CD34^+^ was evident in the murine kidneys, indicating possible engraftment of secondary organs. We also examined whether HSPCs could have potentially adopted a BM niche-like profile (26–28). We found that human SLE-derived PDGFRα^+^ and CD105^+^ cells were present and in an increased percentage in the murine BM, kidneys and spleen, in comparison to healthy-derived human PB progenitors (Figures **5B, C** and **6A**).

These xenotransplantation experiments indicate that human PB HSPCs mediated extramedullary colonisation with an increased ability of the SLE-derived HSPCs to home and engraft at extramedullary sites such as the kidneys, where they may participate in local pathology through the formation of primitive HSPC colonies (Figure **6B**) (28).

**Figure 6.**
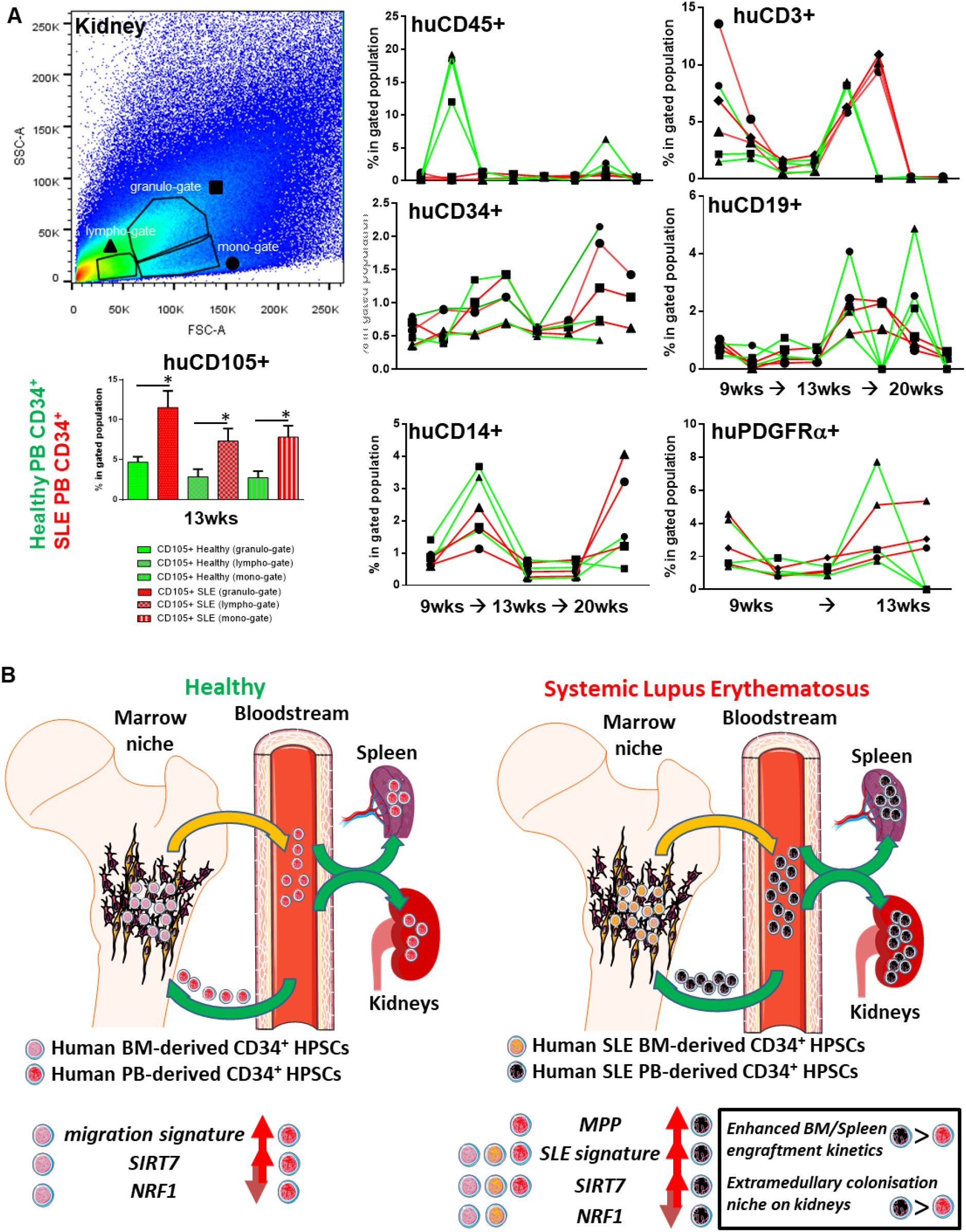
Altered kinetics of SLE CD34^+^ progenitors in the kidney. (A) Mice were sacrificed at designated time intervals and the kidney-derived cells were subjected into immunostaining with antibodies against human-only surface markers, in order to assess human cell presence. Lympho-gates (triangle), granulo-gates (square) and mono-gates (circle) represent different immune cell subpopulations and therefore examined separately, according to their FSC/SSC readout. N=14. Each triangle/dot/circle represents the average value for each time-point and assigned gate. (B) Schematic representation of the kinetics of PB- and BM-derived human CD34^+^ progenitors from healthy and SLE individuals, in relation to NRF1 and SIRT7 differential gene expression as well as their extramedullary potential based on RNA-seq and comparison of SLE and Healthy PB xenotransplantation potential in humanised mice.

### Increased frequency of MPPs in the peripheral blood of SLE patients, with increased extramedullary colonisation in humanised mice

In order to assess whether HSCs/MPPs numbers (found within the CD34^+^ cell population) differ in the periphery of SLE patients, PBMCs were subjected to immunostaining against the surface markers CD34, CD38, CD45RA, CD90 and CD49f, along with the cell viability dye 7AAD. Parallel gating indicated that PB-derived from SLE patients showed an increased frequency in HSC and MPP cell populations when compared to Healthy PB (Figure **7A**). On gated CD34^+^CD38^-^ populations, SLE MPPs were significantly more in BM and statistically significant more in PB (over a two-fold increase) when compared to healthy controls [Figure **7B** and as we have shown recently concerning human healthy and SLE BM CD34^+^ transcriptomes (11)]. No statistical difference was observed in CD34^+^CD38^+^ cell populations (Supplementary Figure **4A**). In addition, disease severity showed to be linked to a two-fold MPP increase in SLE PB (Figure **7B**).

**Figure 7.**
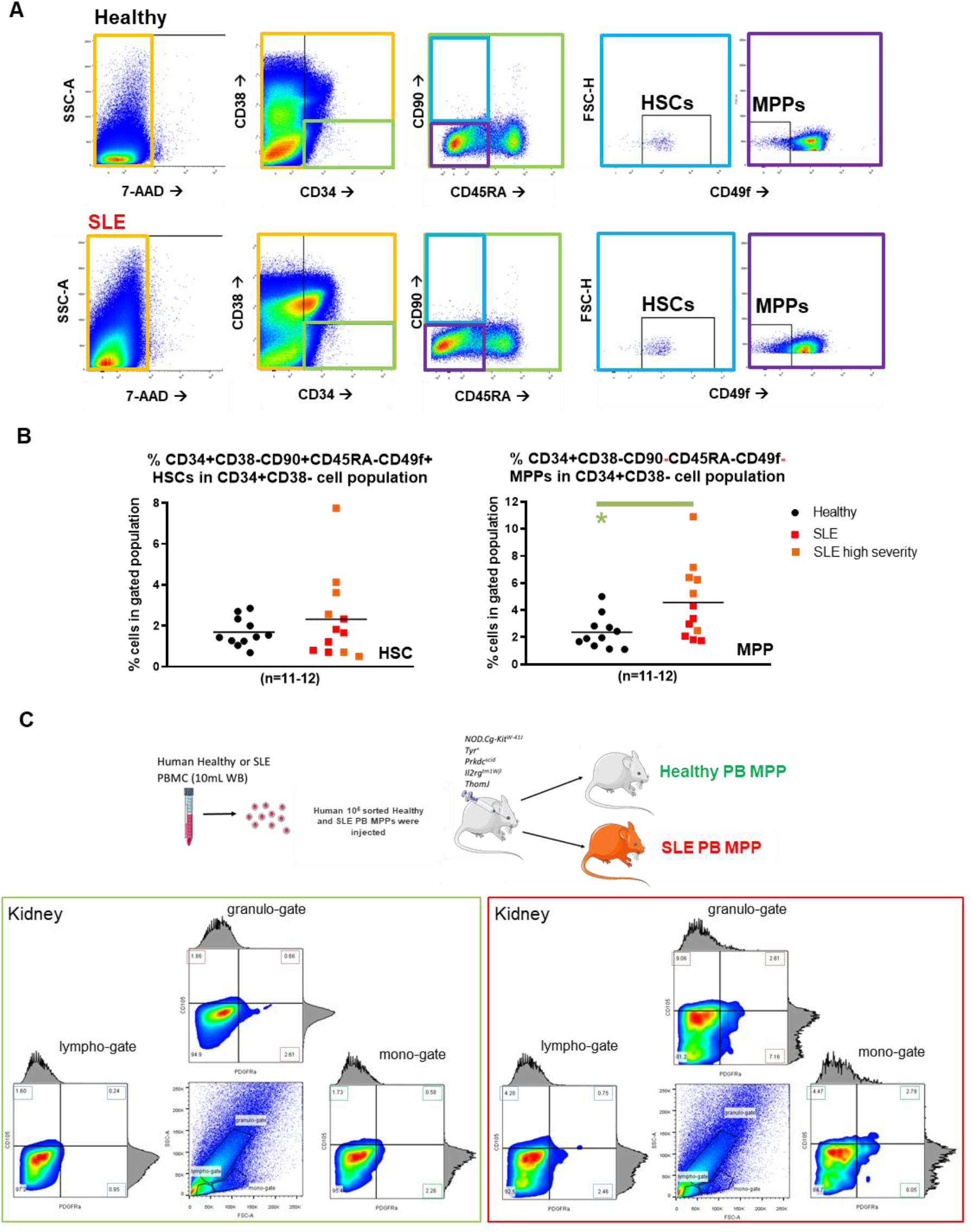
Peripheral blood MPP number is elevated in SLE and migrate preferentially to the kidneys. (A) Gating strategy for obtaining human HSC and MPP cell populations from PBMC from healthy (n=11) and SLE (n=12) individuals. These representative graphs from a total of gated live cell population (7AAD, orange gate) was initially gated for CD34^+^CD38^-^ progenitor pool (green gate), which was further examined for CD45RA^-^CD90^+^CD49f^+^ HSCs (light blue gate) and CD45RA^-^CD90^-^CD49f^-^ MPPs (purple gate). (B) X-Y graphs showing the percentage of HSCs and MPPs in PBMC-gated and CD34^+^CD38^-^-gated cell populations (and mean values). There are statistically more MPPs in the CD34^+^CD38^-^-gated cell populations that is linked to disease severity (orange squares, p<0.03, Student’s T-test). A similar trend was observed in HSCs but was not linked to disease severity (orange squares, p=0.35, Student’s T-test). (C) Mice were sacrificed at 5 weeks and kidney-derived cells were subjected into immunostaining with antibodies against human-only surface markers for CD105 and PDGFRα, in order to assess human cell presence. Lympho-gates, granulo-gates and mono-gates represent different immune cell subpopulations and therefore examined separately, according to their FSC/SSC readout. N=4.

In light of this, SLE or Healthy MPPs were also injected into each of four 2-month old NBSGW mice, and assessed for colonisation potential in the BM, spleen and kidneys, 5 weeks later (one human sample per mouse, Figure **7C** and Supplementary Figure **4B**). FACS analysis indicated that MPPs colonise neither the bone marrow nor the spleen (data not shown). In contrast, only the kidneys were occupied by human MPP-derived CD105^+^PDGFRα^+^ cells, with SLE-derived cells showing an increased presence, when compared to Healthy-MPP-derived cells.

## DISCUSSION

Dysregulation of HSPC activity in the BM has been reported in several chronic inflammatory diseases, including inflammatory bowel disease, atherosclerosis and in SLE. Here, we investigated the potential gene expression differences that involve enhanced migration between PB and BM CD34^+^ HSPCs from both Healthy and SLE patients.

We confirm that PB CD34^+^ HSPCs are distinct from BM CD34^+^ HSPCs as previously shown in a study relying on surface marker intensity alone (29). We also show that the transcriptional landscape is vastly different between BM and PB HSPCs, likely underlying altered outcomes concerning extramedullary colonisation and engraftment success (30). The authors of the latter study also pinpointed, as we did, to the importance of *E2F* TF family gene expression in cell cycle progression. This could be why we see an increase in the number of MPPs in the SLE setting, as confirmed in an SLE murine model in our laboratory (11) (and recent unpublished observations). In fact, what we observed was that PB HSPCs have decreased oxidative phosphorylation and cell cycle signatures as compared to BM HSPCS, indicating a more activated haematopoiesis along with a reduced self-renewal potential (31).

In the SLE setting, PB HSPCs have a gene expression signature that could exacerbate inflammatory responses at local tissues such as the kidneys and other target organs in SLE such as joint and skin. This could be because a number of relevant transcriptional pathways is altered in PB HSPCs, in comparison to BM HSPCs. Of interest, the TLR-7/TNF-alpha/IFN-gamma axis that has been shown recently to be involved in extramedullary damage in murine SLE was highly upregulated in our human PB HSPCs, and especially in SLE PB HSPCs, when compared to BM HSPCs (data not shown) (32). In humanised mice, SLE-PB HSPCs were more prone to colonise extramedullary organs such as spleen or kidneys but not the BM [possibly due to decreased oxidative phosphorylation and self-renewal capabilities (31)], when compared to Healthy-PB HSPCs. SLE-PB CD34^+^ and MPP progenitors may participate in local inflammatory reactions in the periphery due to their increased engraftment/homing potential and altered Th activation (33) while could be promoting an aberrant haematopoietic “niche” (34, 35).

Of particular interest is *NRF1* TF, a master mitochondrial regulator (36), which showed a marked decrease in SLE PB, in comparison to SLE BM (but not healthy PB, data not shown). NRF1 is implicated in pathways related to mitochondrial biogenesis and systemic chronic inflammation (37). A recent publication indicated that SIRT7, a nutrient-sensing protein and a histone deacetylase, when bound to NRF1, is able to alter essential HSC functions related to ageing, possibly *via* mitochondrial function(15) and/or repopulation potential *via* invoking genome stability (when located to the nucleus) (13, 38). We did observe a multi-fold increase in *SIRT7* gene expression, in PB-derived cells, in comparison to BM-derived cells, and a global decrease in oxidative phosphorylation gene expression (Figures 5D and 4B, respectively). These events could be linked to a reduced HSC quiescence with altered regenerative and bone-marrow reconstitution capabilities (13, 14, 23, 24) and, as far as SLE is concerned, direct multi-organ autoimmune inflammation (39) through extramedullary colonisation. Our results corroborate those of a recent study where *E2F1* was identified as one of the susceptibility loci in SLE, directly involving *NRF1* in Asian and European human populations (data not shown) (40). Of note, *SIRT7* and *NRF1* genes were inversely, but not differentially, expressed in Healthy PB vs BM CD34^+^ HSPCs.

A recent study pinpointed that *SIRT1* is involved in recurring infections in SLE patients (41, 42); SIRT7 has the ability to bind directly to SIRT1, possibly being involved in lupus nephritis (43). At the time of the drafting of this paper, Alexander Kaiser *et al*. reported that SIRT7 levels influence remission responses after HSPC transplantation in myeloid leukaemia (44). Indeed, ongoing experiments will focus on investigating if this protein complex confers substantial changes to HSC’s reconstitution capability and other properties such as multiorgan extramedullary colonisation conferring organ-specific SLE tissue injury (model proposed for this study, Figure **6B**). To our knowledge, this is the first study that explores the transcriptome of human non-mobilised circulating HSPCs in the context of a systemic autoimmune disease.

We recognise that an important limitation of the current study is that the expression profile of bulk CD34^+^ HSPCs cannot disentangle minor shifts in the transcriptome of progenitor subpopulations. Yet, this study does pave the way for revealing clinically important transcriptomic differences and migration kinetics between niche and patrolling HSPCs in humans; these findings need to be investigated further using single-cell mRNA-seq technology.

In summary, we show here that the human PB CD34^+^ HSPC transcriptome differs substantially from that of human BM CD34^+^ HSPC, leading to altered engraftment and reconstitution capabilities. These findings underscore the need for employing non-mobilised PB CD34^+^ cells for haematopoietic stem cell therapy (HSCT) regimes, since they may be more beneficial in some autoimmune disease treatments, including SLE, as reported recently (33). These data also shed light to a pathway that involves TF *NRF1* and histone deacetylase *SIRT7* in SLE. Finally, we also demonstrate for the first time that in an autoimmune/inflammatory disease, human HSPCs are not only activated and circulating, but they are also able to migrate and survive in places other than the BM such as the spleen, a site of peripheral immune responses in SLE, as well as at sites of tissue damage, such as kidneys.

## METHODS

### Animals

The NBSGW humanised mouse strain was purchased from Jackson Laboratories (JAX Stock No. 026622) and maintained as homozygotes (*NOD.Cg-Kit^W-41J^ Tyr^+^ Prkdc^scid^Il2rg^tm1Wjl^/ThomJ*). All animals used were 2-3 months of age upon the time of xenotransplantation experiments and fed chow diet in germ-free housing conditions. All mouse animal work has been approved by the BRFAA ethics committee and the Attica Veterinary Department (758634/22-11/2019).

### Isolation of human CD34^+^ progenitors from BM and PB

Human BM aspirate and PB (10 ml each) were collected in EDTA-coated tubes from healthy and SLE patients and subjected to density gradient centrifugation using Histopaque-1077 (Sigma-Aldrich). Briefly, blood was diluted 1:2 with PBS (for PB) and 1:3 PBS (for BM) and carefully layered over Histopaque medium. Tubes were centrifuged at 400 g for 30 minutes (no break) at room temperature. White blood cell layer was carefully collected, and cells were washed with PBS, and treated with red-blood cell lysis buffer (Biolegend, cat no. 420301) prior to magnetic bead separation in order to obtain a minimum of 90% CD34^+^-enriched cell population (human Diamond CD34 isolation kit, Miltenyi Biotec, Bergisch Gladbach, Germany 130-094-531).

For the xenotransplantation experiments in the humanised mice, 2.5×10^5^ human PB CD34^+^ cells were injected retro-orbitally with 100 μl cell suspension into 2-3 months old NBSGW mice. Human conjugated antibodies used for assessing engraftment and differentiation derived from Biolegend (1:100 dilution); huCD45-Brv421 (368522), huCD19-510Br (302241), huCD3-PE (300308), huCD14-FITC (325604), CD34-APC/Cy7 (343514), CD11b-PercP/Cy5.5 (301327), huCD31-PercP/Cy5.5 (303132), huCD73-APC (344005), huCD105-PE (323205) and huPDGFRα-PE/Cy7 (323508).

### Flow cytometry analysis and human HSC and MPP isolation

Freshly isolated human CD34^+^ and peripheral blood mononuclear cells (PBMC) (2×10^6^ cells/sample) were directly immunolabelled with PE anti-human CD34 (clone 561, 1:200), FITC anti-human CD45RA (clone HI100, 1:200), APC anti-human CD38 (clone HB-7, 1:200), PE/Cy7 anti-human CD90 (clone 5E10, 1:200), Brilliant violet 421 anti-human/mouse CD49f (clone GoH3, 1:100) and 7-AAD viability staining solution (1:300), all from Biolegend, in 5% FBS in 1X PBS for 30’ at 4°C. Cells were incubated with the antibodies and viability staining for 30’ at room temperature before washing twice with 1X PBS. Cells were then resuspended in 5% FBS and passed through a 70μm filter pore tip ensuring single cell-only passage for flow cytometric analysis. Flow Cytometry data analysis was performed using FlowJo™ V10.

### NGS

Human CD34^+^ cells were lysed and RNA was extracted using Qiagen’s Nucleospin RNA XS kit, according to the manufacturer’s protocol. mRNA-seq experiments were carried out in the Greek Genome Center (GGC) of Biomedical Research Foundation of the Academy of Athens (BRFAA). mRNA-seq libraries were prepared with the Illumina TruSeq RNA v2 kit using 20-120 ng of total RNA (excluding duplicate values). Libraries were checked with the Agilent bioanalyzer DNA1000 chip, quantitated with the Qubit HS spectrophotometric method and pooled in equimolar amounts for Sequencing. 75 bp single-end reads were generated with the Illumina NextSeq500 sequencer.

### NGS data analysis pipeline

FASTQ data were processed using Cutadapt 1.16 with Python 3.6. (command line parameters: -j 1 -a AGATCGGAAGAGC -a GGGGGGGGGGGGGGGGGGGGGGGGGGGGGGGGGGGGGGGGGGGGGGGGG --output=out1.fq -- error-rate=0.1 --times=1 --overlap=3 --minimum-length=25 --nextseq-trim=30 Concatenate datasets) using the online www.useGalaxy.org platform. BAM files were created using the gapped-read mapper RNA-STAR alignment software with its default values using the GRCh38 primary assembly and the gencode.v29 annotation gtf file (45). BAM files were further sorted using Samtools sort (46). Aligned and sorted BAM files were further analysed on SeqMonk 1.44.0 software. For Gene Ontology (GO) and downstream analysis, Toppgene online tool(47), as well as Cytoscape and ClueGo(48) software were employed.

### Semi-Quantitative PCR

RNA was quantified (by Nanodrop) and retrotranscribed to cDNA (PrimeScript™ RT-PCR Kit - Takara Bio Cat. # RR014A). Real time PCR was performed with KAPA SYBR^®^ FAST qPCR Master Mix (2X) Kit using the following oligonucleotide primers: NRF1 (forward, 5’-CACAGAAAAGGTGCTCAAAGGA-3’; reverse, 5’-CCTGGGTCCATGAAACCCTC-3’), SIRT7 (forward, 5’-CCTGAGCGCGGCCTG-3’; reverse, 5’-GCCTGTGTAGACGACCAAGT-3’), β-actin (forward, 5’-CTCTTCCAGCCTTCCTTCCT-3’; reverse, 5’-AGCACTGTGTTGGCGTACAG-3’). Samples were normalized to β-actin and expression levels were calculated using the 2ΔΔ^-Ct^ method.

### Statistics

Statistical analysis on flow cytometric data was performed using ANOVA T-test using Mann-Whitney or Bonferroni post-hoc test, where appropriate (p<0.05). RNA-seq data statistical analysis was initially performed using p-value (<0.05) and EdgeR for obtaining DE genes. Downstream analysis involved False Discovery Analysis (FDR) based on Benjamini and Hochberg (49).

### Patients & Study approval

Informed consent was obtained from all patients and human controls prior to sample collection (IRB protocol number 10/22-6-2017).

BM aspirates and PB samples were obtained from SLE and gender-matched healthy controls. Patients met the 1999 American College of Rheumatology revised criteria for the classification of SLE (50). All subjects were female apart from two individuals (one SLE and one healthy).

## AUTHOR CONTRIBUTIONS

I.K. designed, conceived and performed the research, analysed the data, and wrote the paper, A.B. and M.G. provided the BM sample data, A.F. performed the initial steps of transcriptome analysis and guided the mRNA-seq data analysis to the correct direction, T.M performed the PCR experiments, T.A. performed the injections in mice and prepared samples for FACS analysis, N.M., I.M., P.V., and D.T.B., advised on several aspects of the research project from guidance to reviewing this paper.

## ACKNOWLEDGEMENTS

We thank Ms Theodora Togia for her excellent lab management and Dr. Anastasia Apostolidou for handling the FACS facility and Dr. Ioannis Vatsellas for RNA-seq processing at the Hellenic Genome Centre in the BRFAA. We also thank M.D. Stavros Doumas for his most insightful comments. Special thanks to Dr Stavroula Giannouli, Dr Antigoni Pieta, M.D. Dionisios Nikolopoulos and M.D. Noemin Kapsala for their help with human sample collection.

