## Supplementary material for "Patrolling human SLE haematopoietic progenitors demonstrate enhanced extramedullary colonisation; implications for peripheral tissue injury": Supplmental Figures 1-4

#### A Healthy vs SLE BM CD34$^{+}$ - CLINVAR (% genes per group)

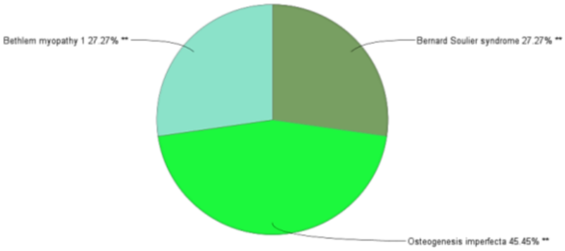


#### B REACTOME Reactions/Pathways (%terms per group)

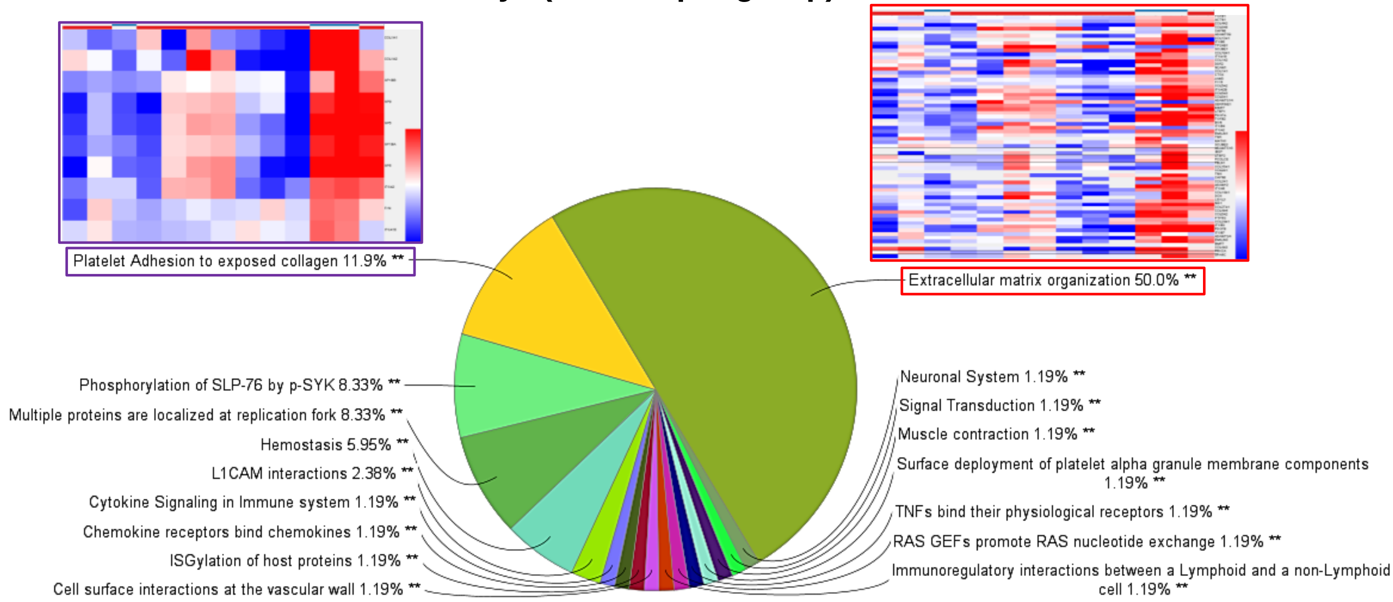


#### C KEGG (%terms per group)

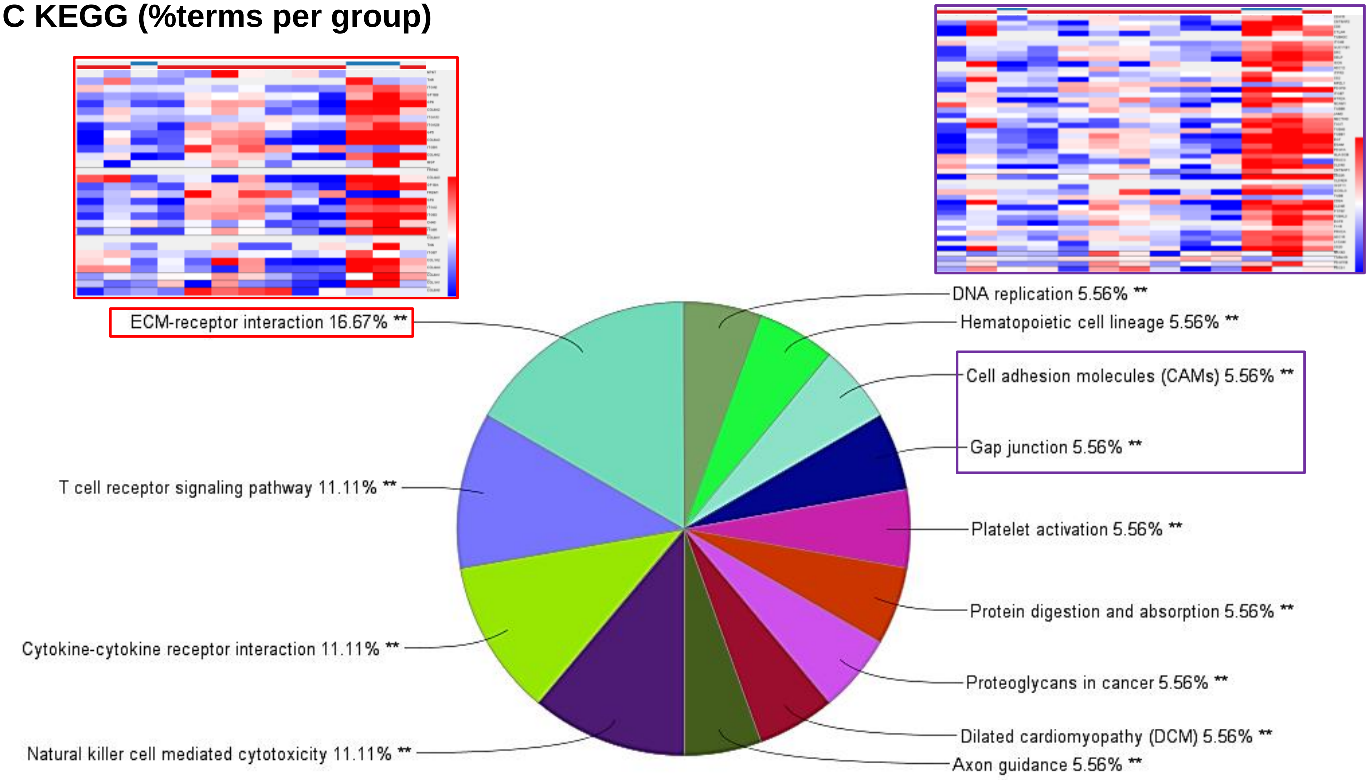

#### Supplementary Figure 2

**Kokkinopoulos *et al.***

##### A CLINVAR and KEGG (SLE BM vs SLE PB)

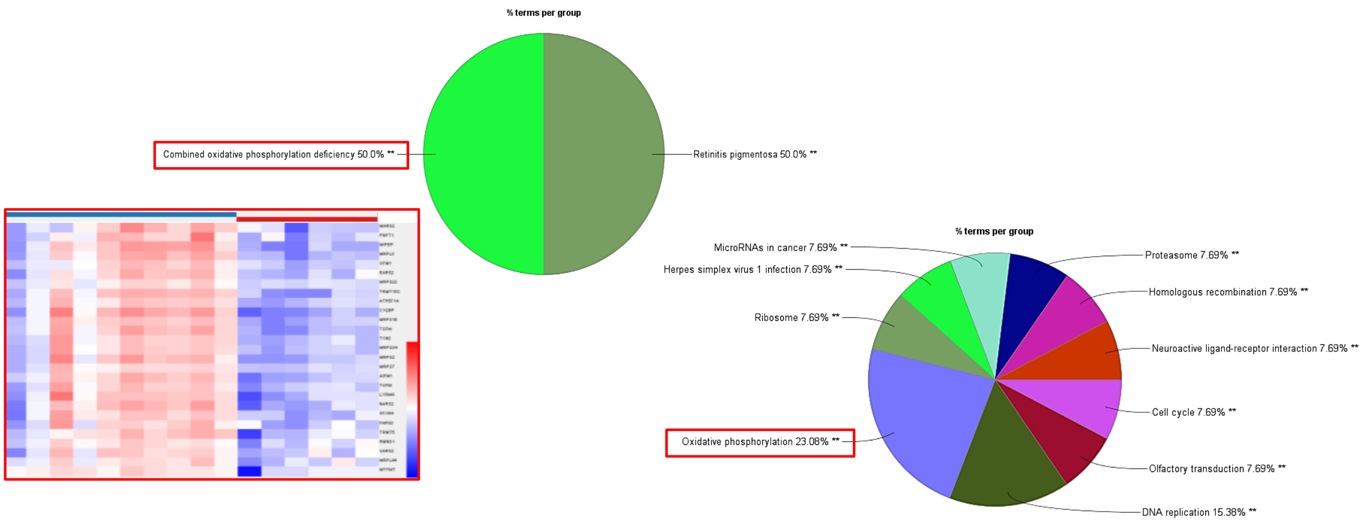


##### B CLIVAR and KEGG (Healthy BM vs Healthy PB)

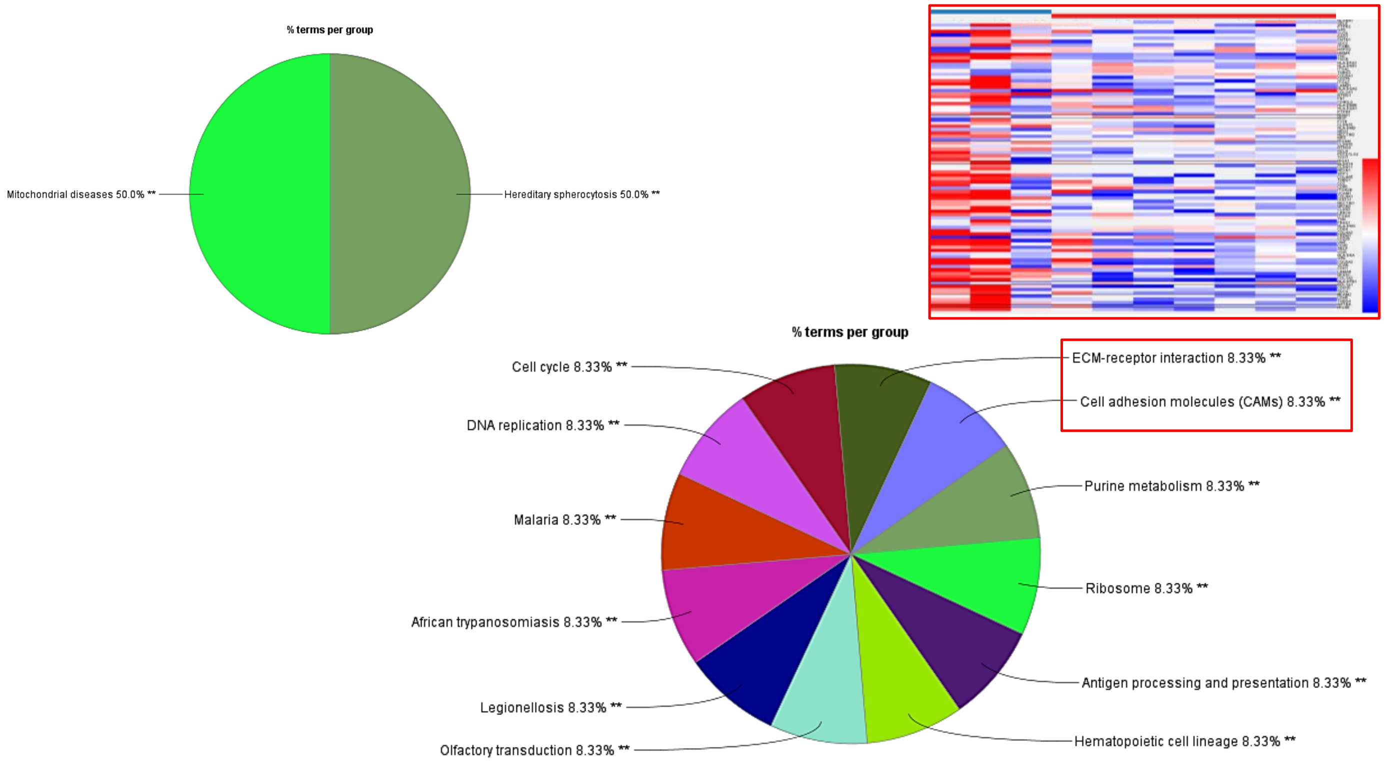


**C**

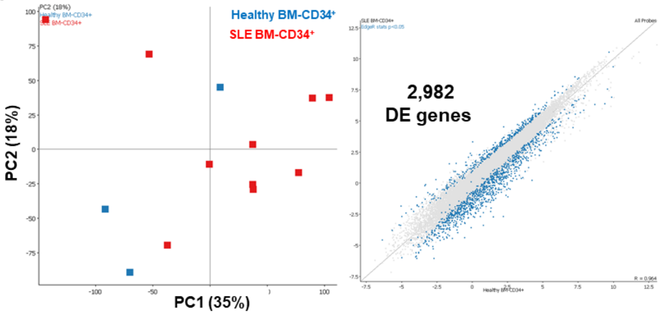


**D**

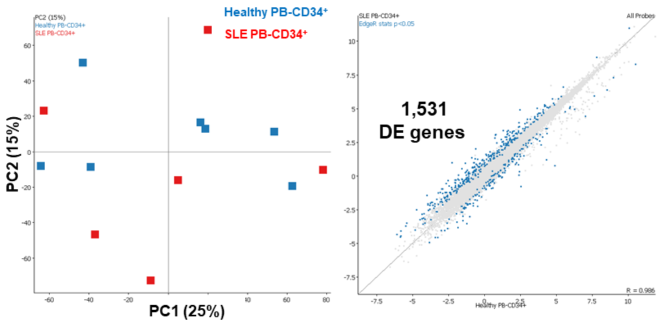

### Supplementary Figure 4

Kokkinopoulos *et al.*

## A

SLE PB vs SLE BM

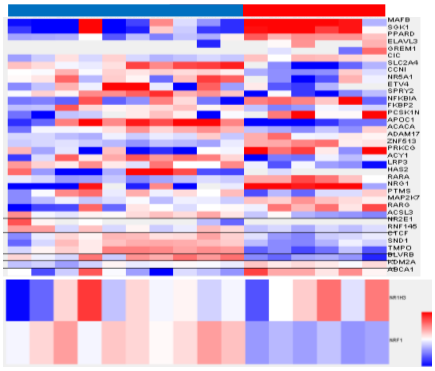

## C

| BM SLE vs Healthy | Hit Count in Query List | q-value FDR B&H | Hit Count in Genome |
|---|---|---|---|
| CAGGTG_V$E12_Q6 | 231 | 9.16E-03 | 1956 |
| V$E12_Q6 | 32 | 3.35E-02 | 201 |
| V$FOXO4_02 | 33 | 3.00E-02 | 205 |
| TTGTTT_V$FOXO4_01 | 202 | 9.76E-04 | 1603 |
| TGGAAA_V$NFAT_Q4_01 | 192 | 9.76E-04 | 1497 |
| V$NFAT_Q6 | 31 | 3.41E-02 | 194 |
| CAGCTG_V$AP4_Q5 | 151 | 8.88E-03 | 1192 |
| GGGTGGRR_V$PAX4_03 | 134 | 1.23E-02 | 1068 |
| CTTTGA_V$LEF1_Q2 | 124 | 1.23E-02 | 981 |
| RYTTCCTG_V$ETS2_B | 114 | 9.16E-03 | 868 |
| RTAAACA_V$FREAC2_01 | 97 | 1.61E-02 | 746 |
| GCANCTGNY_V$MYOD_Q6 | 90 | 4.73E-02 | 721 |
| V$MYOD_Q6 | 38 | 2.71E-03 | 200 |
| V$MYOD_Q6_01 | 34 | 1.23E-02 | 195 |
| CTAWWWATA_V$RSRFC4_Q2 | 46 | 1.55E-02 | 298 |
| CATTGTYY_V$SOX9_B1 | 43 | 3.41E-02 | 293 |
| V$GFI1_01 | 42 | 9.76E-04 | 219 |
| TGATTTRY_V$GFI1_01 | 38 | 2.76E-02 | 244 |
| V$AML_Q6 | 41 | 9.76E-04 | 210 |
| RACCACAR_V$AML_Q6 | 35 | 1.23E-02 | 205 |
| V$GATA1_05 | 39 | 1.00E-02 | 229 |
| V$GATA1_03 | 34 | 1.61E-02 | 203 |
| V$TATA_C | 38 | 3.35E-02 | 248 |
| V$SRF_Q6 | 37 | 9.16E-03 | 211 |
| CCAWWNAAGG_V$SRF_Q4 | 17 | 1.23E-02 | 74 |
| V$SRF_Q5_01 | 33 | 1.55E-02 | 194 |
| V$SRF_C | 31 | 1.68E-02 | 181 |
| V$LYF1_01 | 34 | 4.29E-02 | 222 |
| V$HNF3B_01 | 33 | 9.16E-03 | 179 |
| V$MEF2_02 | 33 | 1.68E-02 | 197 |
| V$MEF2_03 | 33 | 1.79E-02 | 198 |
| V$RSRFC4_01 | 33 | 3.35E-02 | 208 |
| V$E2F1_Q4_01 | 32 | 1.23E-02 | 181 |
| V$RSRFC4_Q2 | 32 | 1.23E-02 | 182 |
| V$E2F_Q3_01 | 32 | 1.55E-02 | 186 |
| V$E2F_Q6 | 30 | 3.18E-02 | 182 |
| V$E2F_Q4 | 30 | 3.35E-02 | 184 |
| V$E2F_Q4_01 | 30 | 3.35E-02 | 185 |
| V$OCT1_04 | 32 | 3.35E-02 | 200 |
| V$PBX1_01 | 32 | 3.35E-02 | 201 |
| V$STAT5B_01 | 32 | 3.41E-02 | 202 |
| V$GATA6_01 | 32 | 4.36E-02 | 206 |
| V$NFKB_Q6_01 | 30 | 3.65E-02 | 187 |
| V$FOXD3_01 | 29 | 1.61E-02 | 164 |
| V$HNF3_Q6 | 27 | 3.35E-02 | 160 |
| V$FOXJ2_01 | 26 | 3.35E-02 | 153 |
| SGCGSSAAA_V$E2F1DP2_01 | 25 | 1.34E-02 | 131 |
| V$AP2REP_01 | 25 | 2.76E-02 | 141 |
| V$PAX6_01 | 18 | 1.23E-02 | 79 |

PB SLE vs Healthy

NO HITS

| BM SLE vs Healthy | Hit Count in Query List | q-value FDR B&H | Hit Count in Genome |
|---|---|---|---|
| AACTTT_UNKNOWN | 202 | 1.55E-04 | 1531 |
| CTGCAGY_UNKNOWN | 84 | 1.03E-02 | 607 |
| TGACATY_UNKNOWN | 76 | 9.73E-03 | 534 |
| YNGTTNNNATT_UNKNOWN | 47 | 9.16E-03 | 286 |
| GTGGGTGK_UNKNOWN | 40 | 9.16E-03 | 234 |
| YYCATTCAWW_UNKNOWN | 28 | 1.68E-02 | 158 |
| RTTTNNNYTGGM_UNKNOWN | 22 | 4.99E-02 | 128 |
| RYAAAKNNNNNNTTGW_UNKNC | 17 | 1.03E-02 | 71 |

## B

Healthy PB vs Healthy BM

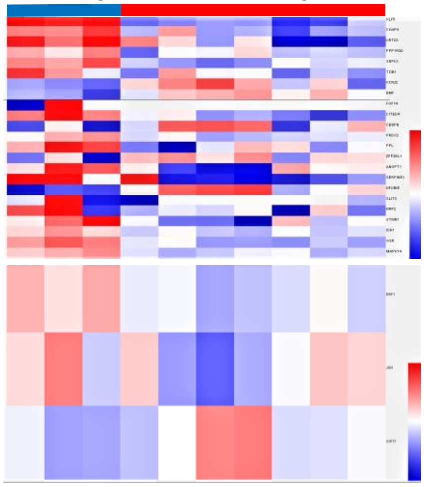

## D

Healthy BM vs SLE BM

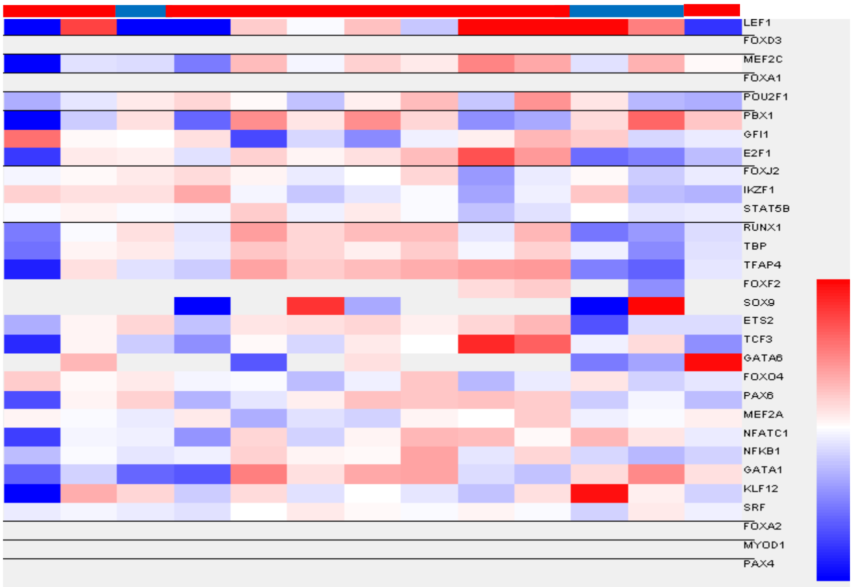


## E

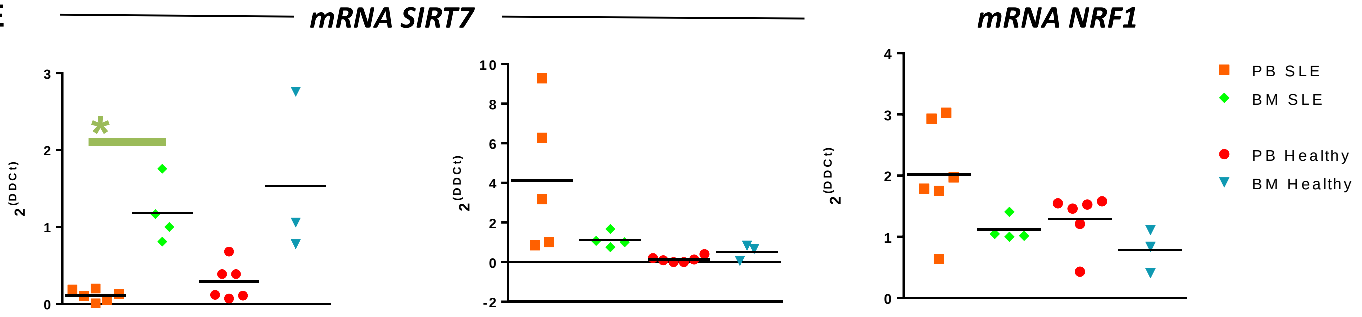

**Supplementary Figure 4** **Kokkinopoulos *et al.***

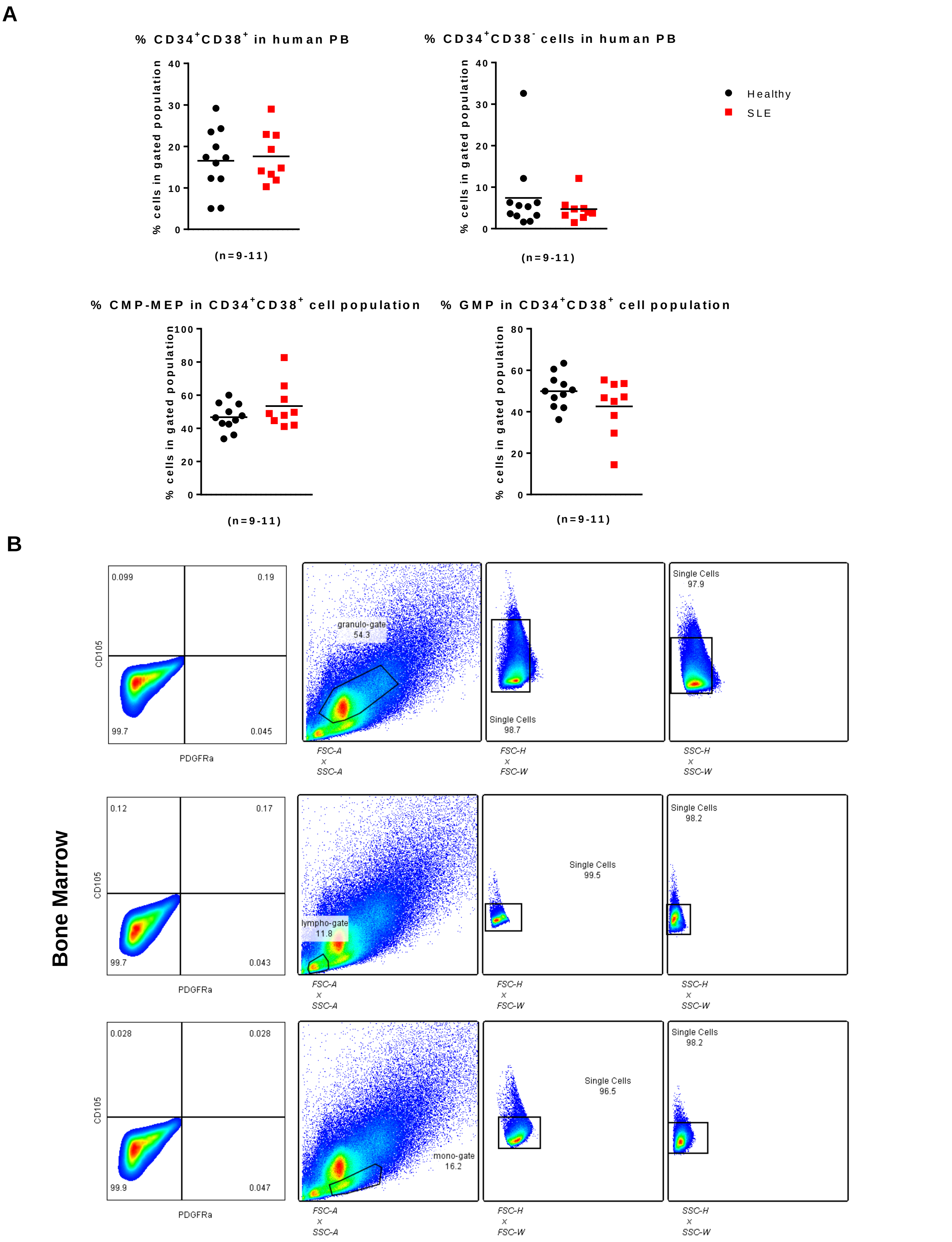

### Supplementary Figure Legends

**Supplementary Figure 1**. (**A**) CLIVAR analysis of DE genes between Healthy BM-derived (Blue) and SLE BM (red) $CD34^+$ progenitors. Analysis were performed with a $p<0.05$ with an FDR correction. (**B**) REACTOME Reactions/Pathways analysis of DE genes between Healthy BM and SLE BM $CD34^+$ progenitors. All analyses were performed with a $p<0.05$ with an FDR correction. (**C**) KEGG Reactions analysis of DE genes between Healthy BM and SLE BM $CD34^+$ progenitors. Analysis were performed with a $p<0.05$ with an FDR correction. Analyses were performed with a $p<0.05$ with an FDR (Bonferroni and Heidelberg) correction. *$p<0.05$, **$p<0.01$.

**Supplementary Figure 2**. (**A**) CLINVAR analysis in SLE PB vs BM and (**B**) Healthy PB vs BM. (**C, D**) PCA analysis and scatter plot of Healthy BM vs SLE BM progenitors and Healthy PB vs SLE PB progenitors revealed an inability to cluster the two circulating progenitor populations, based on disease status. EdgeR analysis indicated 2,982 DE genes between Healthy and SLE BM-derived $CD34^+$ progenitors, while in PB-derived samples, 1,531 DE genes were detected between Healthy and SLE $CD34^+$ progenitors. Analyses were performed with a $p<0.05$ with an FDR (Bonferroni and Heidelberg) correction. *$p<0.05$, **$p<0.01$.

**Supplementary Figure 3**. (**A**) Heatmaps of *NRF1* and *NR1H3* expression in SLE PB (red) and SLE BM (healthy), along with the DE genes that are affected by *NRF1* expression. (**B**) Heatmaps of *NRF1* and *NR1H3* expression in Healthy PB (red) and Healthy BM (blue), along with the DE genes that are affected by *CEBPB*. (**C**) Table indicating the predicted TF binding sites (Toppgene software) based on the DE gene input lists for BM SLE vs Healthy, PB SLE vs Healthy and BM SLE vs Healthy. (**D**) Heatmaps of expression for TFs of interest in Healthy BM (blue) and SLE BM (red). All analyses were performed with a $p<0.05$ with an FDR (Bonferroni and Heidelberg) correction. (**E**) Real-time PCR graphs for *NRF1 variants (ENST00000353868.5, ENST00000393232.5, ENST00000311967.6, ENST00000223190.8)* and *SIRT7 variants (ENST00000572902.5, SIRT7-210 and ENST00000328666.11, SIRT7-201)* in all four groups. The expression of *NRF1* was not statistically different in any of the four groups. Some *SIRT7* variants were downregulated significantly in SLE PB in comparison to SLE BM $CD34^+$ progenitors. One-way ANOVA with Bonferroni's post-hoc test, $p<0.05$, n=3-6.

**Supplementary Figure 4**. (**A**) Peripheral blood CMP-MEP and GMP numbers remain the same in PB Healthy vs SLE. X-Y graphs showing the percentage of $CD34^+CD38^+$, $CD34^+CD38^-$, CMP-MEP and GMP in PBMC-gated and $CD34^+CD38^-$-gated cell populations. n=9-11, Student's T-test. (**B**) Representative diagrams of the gating strategy employed for examining human MPP-derived cell populations in the bone marrow of humanised mice. A similar strategy was used upon examination of kidney and spleen tissue.
